# Integrating Gene Expression Analysis and Ecophysiological Responses to Water Deficit in Leaves of Tomato Plants

**DOI:** 10.1101/2024.07.05.602262

**Authors:** G Bortolami, T de Werk, M Larter, A Thonglim, B Mueller-Roeber, S. Balazadeh, F. Lens

**Affiliations:** Naturalis Biodiversity Center, Research Group Functional Traits, PO Box 9517, 2300 RA Leiden, The Netherlands; Plant Ecology Research Laboratory, School of Architecture, Civil and Environmental Engineering, 1015, Lausanne, Switzerland; University of Potsdam, Institute of Biochemistry and Biology, Karl-Liebknecht-Straße 24-25, Haus 20, 14476 Potsdam, Germany; Max-Planck Institute of Molecular Plant Physiology, Am Mühlenberg 1, 14476 Potsdam, Germany; BIOGECO, INRA, Université de Bordeaux, 33615 Pessac, France; Leiden University, Institute Biology Leiden, Sylvius Laboratory, Sylviusweg 72, 2333 BE Leiden, The Netherlands

**Keywords:** Water deficit, crops, ecophysiology, gene expression, hydraulics, embolism, ABA-dependent, ABA-independent, transcription factors

## Abstract

Soil water deficit (WD) is one of the most important abiotic stresses affecting plant survival and crop yield. Despite its economic relevance, many gaps remain in our understanding of how crops respond to WD, especially concerning the synergistic coordination of molecular and ecophysiological adaptations delaying plant damage and mortality. In this study, we investigated the gene expression imposed by a progressive WD and combined it with measurements pointing to key ecophysiological thresholds in leaves of tomato plants. We uncovered the transcriptomic changes in mature leaves at four stages defined by physiological markers relating to different WD intensities: partial stomatal closure, complete stomatal closure, after leaf wilting, and beginning of embolism development in the veins. By identifying key transcription factors (TFs) across these progressively worsening WD stages, we investigated the timing and impact of ABA-(in)dependent gene regulatory pathways during WD. In addition, we compared the transcriptome in young developing versus mature leaves and explored the physiological mechanisms that may explain the higher tolerance to dehydration in younger leaves. By correlating the transcriptomic changes to precise ecophysiological measurements, the combined dataset will serve as a framework for future studies comparing leaf molecular and physiological responses to WD at specific intensities.

**Highlight:** Integrated ecophysiological and gene expression analyses identify key mechanisms underlying the different thresholds of tomato responses to water deficit

## Introduction

The environmental conditions, encompassing all abiotic and biotic factors, define the niche boundaries where a plant species can actively adapt and grow (Chase & Leibold, 2003; Treurnicht *et al.*, 2020). Any deviation in these conditions from the optimal niche range leads to plant stress and a reduction in growth and yield (Bailey-Serres *et al.*, 2019; Zhang *et al.*, 2022). In the present age of anthropogenic climate change, it is evident that global hydrological patterns are swiftly evolving, leading to diminished precipitation in numerous geographical areas and making them less suitable for agriculture (Greve *et al.*, 2014, 2017, 2018; Wartenburger *et al.*, 2017; Xu *et al.*, 2019). In a world where plant growth will become more demanding, there is an urgent need to focus more on mechanisms underlying drought responses in crops to safeguard food production (Pingali, 2012; Ray *et al.*, 2013). Tomato (*Solanum lycopersicum* L.) is one of the major crop species cultivated in the Mediterranean area where droughts are projected to become more intense and more frequent (Guerreiro *et al.*, 2018; Raymond *et al.*, 2018; Lionello & Scarascia, 2020). Previous research investigating tomato response to water deficit (WD) is scattered and does not provide a complete picture of plants’ sensibility to stress. Available drought studies in tomato either focused on only one single WD intensity (Gong *et al.*, 2010; Iovieno *et al.*, 2016; Mishra *et al.*, 2016; Zhou *et al.*, 2019; Diouf *et al.*, 2020; Bian *et al.*, 2021; Pirona *et al.*, 2023), used osmotic stress or dehydration as a substitute for diminishing soil water availability (Dong *et al.*, 2023; Pirona *et al.*, 2023), examined the WD response only during the seedling stage (Gong *et al.*, 2010; Zhou *et al.*, 2019; Pirona *et al.*, 2023), or investigated the physio- anatomical drivers of xylem embolism development (Lamarque *et al.*, 2020; Harrison Day *et al.*, 2022; Haverroth *et al.*, 2024).

Traditionally, when plant responses to WD are studied, two main approaches are applied to investigate the sequence of events leading to drought-induced plant mortality. The first set of studies focuses on the ecophysiological and/or anatomical approach, looking at modifications of the long-distance water transport pathway and/or changes in xylem anatomical traits imposed by different levels of soil water content (Lens *et al.*, 2011; Martin- StPaul *et al.*, 2017; Corso *et al.*, 2020; Lamarque *et al.*, 2023; Lens *et al.*, 2023). A second line of publications is molecular-oriented and investigates the finer-scale changes in the gene, protein, and chemical chain of reactions during different stages of WD (Hummel *et al.*, 2010; Taiz *et al.*, 2024). Surprisingly, these two scientific disciplines are typically separated from each other and focus on different plant groups: the hydraulic-anatomical approach deals almost exclusively with tree species, studying the sequence of events triggering the forest decline (Choat *et al.*, 2018; Mantova *et al.*, 2022), while the molecular approach mainly focuses on non-woody model species (one above all, *Arabidopsis thaliana*) subjected to induced stress under controlled growth chamber conditions (Lamesch *et al.*, 2012; Singh *et al.*, 2022). Cross-pollination between the two disciplines is only sporadic, mostly limited to the quantification of the expression of key genes during the ecophysiological monitoring of species subjected to WD (Zhang *et al.*, 2020; Thonglim *et al.*, 2023), or constrained to the measurements of leaf water potential (Ψ_leaf_) and gas exchange when investigating gene regulatory pathways imposed by WD (Iovieno *et al.*, 2016; da Silva *et al.*, 2022). Finally, only a few studies performed ecophysiological investigations in plants over- or underexpressing selected genes (Lamarque *et al.*, 2020, Thonglim *et al.*, 2021). Consequently, achieving a deeper comprehension of plant responses to WD demands a more integrated approach, wherein the comparison of gene expression at specific time intervals during WD - based on ecophysiological monitoring - is essential.

Here, we performed a holistic drought study on the leaves of tomato plants. We outlined three key objectives: firstly, we coupled precise ecophysiological monitoring (water potential, gas exchange, stomatal and xylem anatomy, xylem embolism spread, turgor loss point) of fully developed leaves in mature (around 50 days old) tomato plants with quantification of gene expression in leaves at four WD intensities: 1) partial stomatal closure, 2) full stomatal closure, 3) leaf wilting and the start of embolism formation, and 4) the point where embolism starts to exponentially increase corresponding to 12% loss of hydraulic conductivity in the main veins. Secondly, we identified and compared the gene expression at these four physiological thresholds. Thirdly, we focused on a leaf developmental stage-specific response, providing a putative transcriptomic basis for the higher resilience of younger tomato leaves to WD.

## Materials and methods

### Plant material, sampling, and water deficit experimental design

Tomato (*Solanum lycopersicum* cv. MoneyMaker) seeds were germinated on full-strength Murashige-Skoog media containing 3% sucrose (MS 3%). After three weeks, plants with the most developed hypocotyls were transferred into 3L pots in a mixture of potting soil (basis biomix, Lensli® substrates, Bleiswijk, The Netherlands), vermiculite and sand in a ratio of 25:8:2, along with three tablespoons of osmocote fertilizer (ICL Growing Solutions, Geldermal, The Netherlands). At the Institute of Biology Leiden (Leiden University), all potted plants were placed in the same growth chamber at 24°C and 70% relative humidity, with a daily 16-h light and 8-h night cycle at 200 µmol/m^2^s^-1^ photosynthetic active radiation and watered every other day until the beginning of the water deficit experiment. For the RNA-Seq sampling, we divided 50-day-old plants into two groups: a well-watered (WW) control group and a water deficit (WD) group that was fully deprived of water from this point forward. To safeguard against potential plant harm that could impact gene expression analysis while monitoring WD intensity, leaf water potential (Ψ_leaf_) was assessed every other day in a subgroup of plants (three WW and three WD individuals). This was done until critical thresholds were reached, based on already available leaf vulnerability curves linking Ψ_leaf_ with drought-induced embolism formation in leaf veins using a previous batch of plants with equal age (see section “Non-invasive optical determination of embolism vulnerability in leaves”). The monitored plants were not utilized for RNA extraction. The Ψ_leaf_ measurements were conducted using a pressure chamber (Model 1000, PMS Instrument Company, Albany, OR, USA) on a leaflet from a fully matured leaf, following the procedure recommended by Rodriguez-Dominguez *et al.* (2022). When a specific WD intensity was reached (i.e., at 4, 5, 8, and 10 days after water deficit; Fig. 1, Suppl. Fig. S1), a mature leaf from the 6-8^th^ node from the base was used for ecophysiological monitoring, and RNA extraction as detailed in Supplementary Fig. S2.

**Figure 1.**
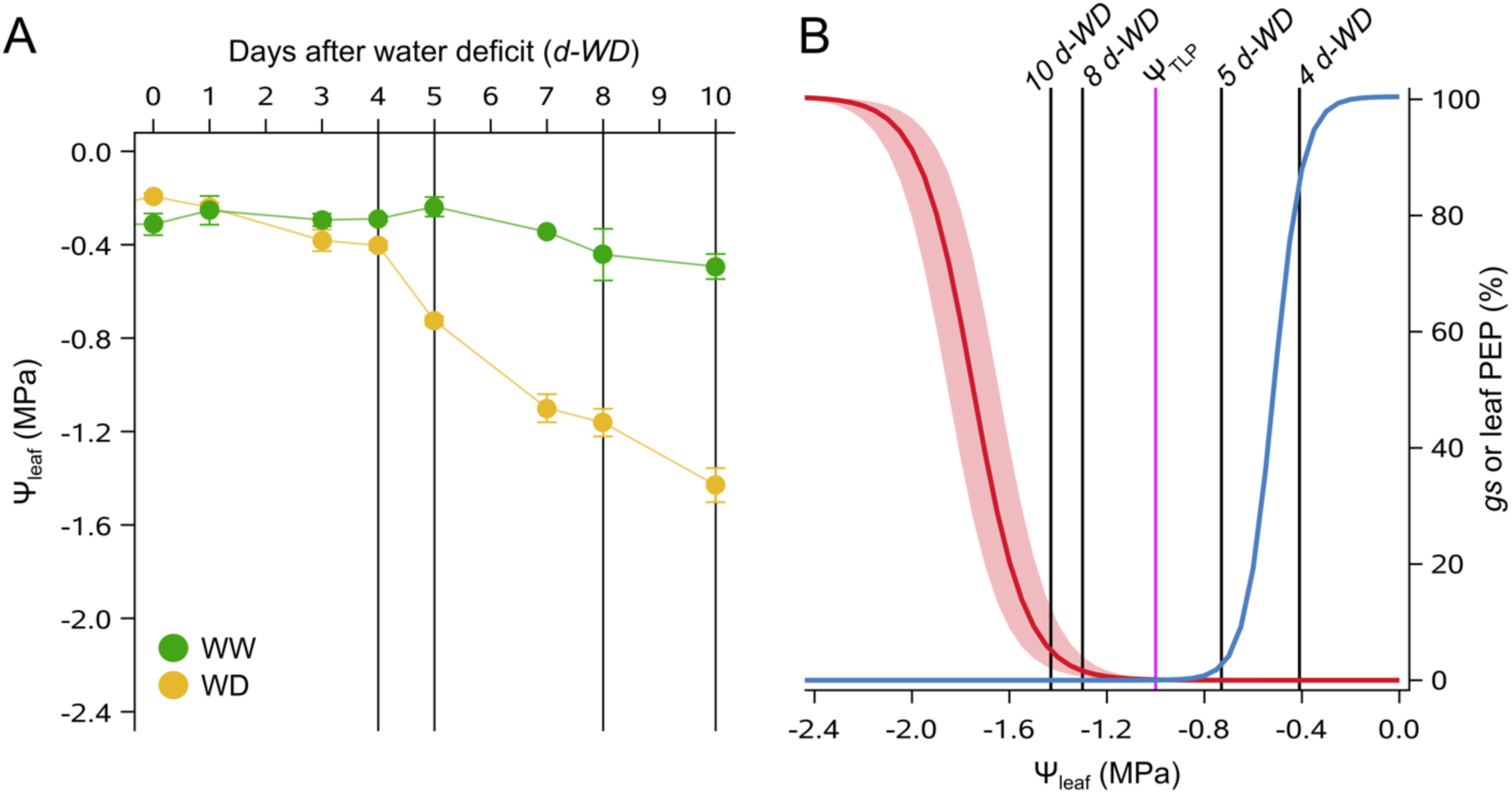
Sampling strategy and physiological thresholds in 50-day-old *S. lycopersicum* cv MoneyMaker plants. **A)** Leaf water potential (Ψ_leaf_) variation over time for well-watered (WW, green circles) and water-deficit (WD, yellow circles) plants during the 10d drought experiment. The four sampling points for RNA extraction (at 4, 5, 8, and 10 days after water deficit, d-WD) are indicated with black vertical lines. Symbols represent averages ± SE (n=3-6 per sampling point). **B)** Stomatal safety margin during Ψ_leaf_ decline, defined by the difference between the best-fitted blue curve for stomata dynamics (*gs*, n=33), and the red vulnerability curve (average ± SE) based on leaf percentage of embolized pixels (leaf PEP, n=8). The purple vertical line indicates the turgor loss point (Ψ_TLP_), and the black vertical lines indicate the four sampling dates for RNA extraction.

### Single-leaflet gas exchange measurements and stomata anatomy

Maximal leaf gas exchange measurements were registered between 9:00 a.m. and 2:00 p.m. on a well-exposed leaflet close to the one used for RNA extraction (Suppl. Fig. S2) using a TARGAS-1 portable photosynthesis system (PP Systems, Amesbury, MA, USA). Tomato plants (n=33, including the ones used for RNA-Seq analyses) were grown in multiple batches from January until November 2022 as described above, and measured between 45 and 60 days after sowing. Optimal conditions of photosynthetic active radiation were set in the cuvette (1500 μmol photons m^-2^ s^-1^). The following parameters were recorded on each selected leaflet: stomatal conductance (*gs*, mmol H_2_O m^-2^ s^-1^), assimilation (*A*, μmol CO_2_ m^-2^ s^-1^), and water use efficiency (*WUE* = *A gs*^-1^, μmol CO_2_ mmol H_2_O^-1^). The same leaflet from four well-watered plants was later stored in 70% ethanol for anatomical observations of stomata density and size. Before these stomatal measurements could take place, the samples were further dehydrated using a series of 70%, 96%, and 100% ethanol, submerged in acetone, and placed in the critical point dryer (Leica EM CPD300, Leica Microsystems, Wetzlar, Germany), mounted on scanning electron microscope (SEM) stubs (12.2x10 mm, Ted Pella Inc., Redding, CA, USA) using carbon adhesive tabs (12 mm diameter, Electron Microscopy Sciences, Hatfield, PA, USA), and coated with a Quorum Q150T S Platinum Palladium sputter coater (Quorum Technologies, Laughton, UK). The abaxial leaf side was photographed with an SEM (JSM-6480 LV, JEOL, Tokyo, Japan) at 5 kV. Stomata were counted in six photos per sample, covering a total area of 1.84 mm^2^, and their length was measured on 36 random stomata per sample.

### Intervessel pit membrane thickness

The thickness of the intervessel pit membranes in tomato leaves was measured in the leaflets xylem of five different plants before the determination of vulnerability to embolism, see below. We detached a leaflet from a node close to the one used for embolism detection, we collected a 1 cm-long section and immediately fixed it in Karnovsky’s fixative. As described in Thonglim *et al.* (2021), the samples were cleaned three times in 0.1 mM cacodylate buffer, then post-fixed with 1% buffered osmium tetroxide, rinsed again with buffer solution, stained with 1% uranyl acetate, and dehydrated in a series of ethanol: 1% uranyl acetate replacement, with increasing concentration of ethanol (30, 50, 70, 96%, and twice in ≥99%). The samples were then infiltrated with Epon 812n (Electron Microscopy Sciences, Hatfield, UK) and placed at 60°C for 48 h in an oven. The Epon blocks were trimmed to a thickness of 2 μm using a rotary microtome with a glass knife. Subsequently, the cross-sections with many vessel–vessel contact areas were cut into ultrathin sections of 90–95 nm using a Leica EM UC7 ultramicrotome with a diamond knife. The sections were dried and mounted on film-coated copper slot grids with Formvar coating (Agar Scientific, Stansted, UK), and post-stained with uranyl acetate and lead citrate. Ultrastructural observations of intervessel pits were performed and photographed using a JEM-1400 Plus TEM (JEOL, Tokyo, Japan) equipped with an 11-megapixel camera (Quemesa, Olympus). The photos were analyzed with ImageJ (Schneider *et al.*, 2012), and the thickness was measured in three different spots of the membrane. We measured the thickness of six to fourteen intervessel pit membranes from five different leaflets, for a total of fifty-two membranes.

### Non-invasive optical determination of embolism vulnerability in leaves

Air embolism formation and propagation in the leaf xylem were measured by adapting an optical technique (Brodribb *et al.*, 2016) monitoring light transmission through the leaf veins. Tomato plants (n=8) were grown in multiple batches from September 2020 until January 2022 as described above and measured between 45 and 60 days after sowing. Plants were moved to the nearby hydraulics laboratory at Naturalis Biodiversity Center, where the soil was gently washed out without damaging the root system. One leaflet (still attached to the plant and in the same position as the one used for RNA-Seq or water potential measurements, Suppl. Fig. S2) was fixed to a scanner (Perfection V850, Seiko Epson Corporation, Suwa, Japan) and scanned every five minutes during one week of dehydration using an automating IT script developed by platform Phenobois Bordeaux, while the stem water potential (Ψ_stem_) was measured at least once a day with the pressure chamber using a bagged leaflet. The resulting images were analyzed with ImageJ software (Schneider *et al.*, 2012) following instructions from http://www.opensourceov.org (last consulted November 2023). The formation of embolisms over time was determined by subtracting the differences in pixels in the 1^st^- to 3^rd^-order veins among the subsequent image scans. Background noise, mainly caused by tissue shrinkage, was removed using mild filters and manual image inspection. Vulnerability curves corresponding to the percentage of embolized pixels (PEP) as a function of Ψ were fitted for every sample based on the following equation (Pammenter and Van der Willigen, 1998):

PEP = 100 / {1 + exp [S/25 x (Ψ-P_50_)]}

where P_50_ is the xylem pressure at which 50% of embolized pixels were observed and S is the slope of the curve at the inflection point.

### Pressure-volume curves and determination of turgor loss point

We determined the leaf turgor loss point (Ψ_TLP_) in four leaflets from two well-watered plants right after the last sampling point using the pressure-volume curve method (Sack *et al.*, 2010). Briefly, plants were watered at pot capacity, and the next day a leaflet from a mature fully developed leaf from the 6-8^th^ node (comparable to the one used for RNA-Seq, Suppl. Fig. S2) was detached and equilibrated inside a sealed plastic bag. The leaf was weighed with a precision scale (Practum224-1S, Sartorius, Göttingen, Germany), its water potential was measured with the pressure chamber, and afterward air-dehydrated in the lab. Starting from the onset of dehydration, its weight and water potential were measured two to three times per day for three days until complete dehydration. Pressure-volume relationships were constructed for every sample by plotting the inverse of leaf water potential (–1/Ψ_leaf_) against the relative water content (RWC), and Ψ_TLP_ estimated as the point of transition between the linear and nonlinear portions of the pressure-volume relationship.

### RNA extraction and library preparation

A leaflet from a mature and fully developed leaf (Suppl. Fig. S2) was taken from the 6-8^th^ node from the base and flash frozen in liquid nitrogen. In the latest sampling date (10 *d-WD*), also young developing leaves were sampled from the topmost (closest to the shoot apical meristem) secondary shoots. All samples were collected between 2 p.m. and 3:30 p.m. In each case, the target leaflets from two plants were combined to form a single biological replicate, and each time point contained two to three biological replicates for the WW and WD batch. Tissues were ground into a fine powder using a Tissuelyzer (QIAGEN), ensuring the samples remained frozen. Total RNA was extracted using TRIzol® Reagent (Thermo Fisher) according to the manufacturer’s instructions. Total RNA purity was assessed using 260/280 and 260/230 nm wavelength absorption ratios, using a NanoDrop™ spectrophotometer (DeNovix DS-11 FX+). Library preparation and paired-end sequencing were performed by BGI Genomics (China). In total, 27 samples had RNA of sufficient quality and quantity to be sequenced (Suppl. Table S1).

### Differential expression and co-expression analyses

Read trimming, alignment of reads to the genome, and mapping of reads to gene models were performed using CLC Genomics Workbench (Qiagen). Reads were aligned to the SL4.00 genome release, using ITAG4.1 gene annotation data (obtained from the solgenomics FTP site: https://solgenomics.net/ftp/tomato_genome/, accessed in June 2023). Automated human readable gene descriptions (AHRD) corresponding to the ITAG4.1 genome were downloaded from the Solgenomics website and added manually (Solgenomics, 2020). Differential expression analyses were performed in R (version 4.3.1), using the DESeq2 package (Love *et al.*, 2014) between WD and WW samples on every sampling date. Correction for the overestimation of effect sizes for lowly expressed genes was done with a log_2_ fold change (LFC) shrink, as described by Zhu *et al.* (2019). Differentially expressed genes were defined as having an LFC difference of at least 1 in the WD plants compared to the control, and a Benjamini-Hochberg FDR adjusted p-value (p_adj_; as given by DESeq2) below 0.05 (Suppl. Table S2). Differential gene regulation between developing and mature leaves was determined using the DESeq2 package, calculating differential regulation on the interaction between leaf age and treatment while controlling separately for both leaf age and treatment. Genes were defined as differentially regulated using a stringent filter (LFC > |2|; *p_adj_* < 0.05, Suppl. Table S3) to account for the increased number of independent variables and reduce the chance for false positives. K-means clustering was performed on Variance Stabilizing Transformation (VST)-normalized counts, as obtained from the raw read counts using DESeq2 in R. Clustering accuracy was validated manually by plotting data grouped by clusters along its three principal components (Suppl. Fig. S3).

### Weighted gene correlation network analysis (WGCNA), co-expression analysis, and identification of water deficit markers

Normalized counts were obtained via a VST from the unaltered read counts using the DESeq2 package. The weighted gene co-expression network analysis (WGCNA) was performed on differentially expressed genes using the WGCNA package in R (version 4.3.1), using a soft power threshold of 20, generating a signed network on the previously normalized counts (chosen as described in Langfelder and Horvath, 2008; Suppl. Fig. S4). Cluster-Trait relationships (using watering regime, sampling date, position of the leaf on the plant, Ψ_leaf_, *gs*, *A*, and leaf PEP as functional traits) were calculated from the cluster eigengenes as described in Langfelder and Horvath (2008). Genes encoding for transcription factors (TFs) were identified from the GO term “DNA-binding transcription factor activity”. TFs of interest were subsequently identified as those that presented an LFC higher than 4 when summed across all time points. The co-expression network was generated in Cytoscape using the CoExpNetViz app (Tzfadia *et al.*, 2016), using previously identified transcription factors of interest as bait genes. Transcription factors of interest were taken as being differentially expressed in WD plants compared to WW plants (LFC > |1|; *p_adj_*-value < 0.05) at any time point, while the networks were generated on all differentially expressed genes irrespective of time point. A stringent filter was applied for significant correlations (p-value ≤ 0.05), with a Pearson’s correlation (r) ≥ |0.95| used to generate the co-expression network. To identify which nodes exerted the most influence on the network - and thus would represent potential transcription factors of interest - we additionally calculated betweenness-centrality between all nodes in Cytoscape. The number of correlated genes for each transcription factor of interest was extracted from the topological overlap matrix, using a weight threshold of 0.4. To visualize the gene expression clusters, z-scores were calculated from the normalized expression values by centering each value on the mean expression and scaling it by the standard deviation of the entire dataset.

## Results and discussion

### Ecophysiological responses of tomato leaves to increasing water deficit

We monitored ecophysiological changes in plants subjected to a 10d period of water deprivation, while simultaneously taking samples for gene expression analyses at specific time points determined by the monitoring (Fig. 1). Under well-watered (WW) conditions where the leaf water potential (Ψ_leaf_) was above -0.4 MPa, the plants showed a leaflet stomatal conductance (*gs*) of 392±18 (average±SE) mmol H_2_O m^-2^ s^-1^ and a CO_2_ assimilation (*A*) of 8.9±0.4 µmol m^-2^ s^-1^. These leaflets showed 154±5 stomata mm^-2^ with a length of 18.35±0.2 µm at the abaxial side and an intervessel pit membrane thickness of 0.42±0.01 μm. As the soil water content decreased, Ψ_leaf_ declined as well (Fig. 1A) and stomata closed (Fig. 1B) at Ψ_leaf_=-0.7 MPa. The turgor loss point in leaves (Ψ_TLP_, the key physiological indicator of a plant’s ability to maintain cell turgidity in leaves, defined when the pressure in- and outside the cells is equal) was identified at -1±0.04 MPa (purple line in Fig. 1B). At a later stage of WD, air embolism appeared and spread in the leaf veins, causing 12% increase of embolized pixels (12% PEP, or P_12_) at -1.41±0.1 MPa, while the average P_50_ and P_88_ were detected at -1.75±0.1 and -2.08±0.14 MPa, respectively (Fig. 1B). The stomatal safety margin (i.e., the xylem pressure range between stomatal closure and the P_50_) corresponded to 1.05 MPa.

To elucidate the molecular processes underlying the observed physiological changes, we extracted RNA at distinct physiological stages for a comparative transcriptomic analysis through RNA sequencing (Fig. 1, 2). To this end, we sampled comparable leaflets from mature leaves for the WW condition and at 4, 5, 8, and 10 days after water deficit (d-WD). These sampling dates corresponded to precise physiological thresholds: at 4d-WD the stomata started to close (*gs_10_*, Ψ_leaf_=-0.41 MPa), at 5d-WD the stomata were completely closed (*gs_90_*, Ψ_leaf_=-0.73 MPa), at 8d-WD the leaves appeared wilted and embolism started to develop in the xylem (4-6% PEP, Ψ_leaf_=-1.3 MPa), and at 10d-DW, P_12_ was reached (Ψ_leaf_=-1.43 MPa). In addition, at 10d-WD, we also sampled young developing leaves closer to the stem apex that showed no wilting despite being under a lower Ψ_leaf_ compared to mature leaves on the same plant (Ψ_youngleaf_=-1.61 versus Ψ_matureleaf_=-1.43 MPa on average).

### Transcriptomic profiling during increasing water deficit

Over 1 billion paired clean reads were successfully mapped across all samples to the SL4.00 genome release, with 93.35% to 95.47% of counted fragments uniquely mapped to genes according to the ITAG4.1 gene annotation release (Suppl. Table S1). A multidimensional scaling plot shows that all the WW samples cluster close together, showing comparable transcriptomic profiles and indicating that our control samples are similar (Fig. 2A). In contrast, a clear progressive separation over Dimension 1 (representing 61% of the observed variance) is seen for WD samples, correlated with progressively increasing WD (Fig. 2A). Similarly, hierarchical clustering analysis of the top 500 most variable genes across all samples clustered all replicates to their respective time points and conditions (Fig. 2B), highlighting the low variability between biological replicates and the overall good quality of the sampling.

**Figure 2.**
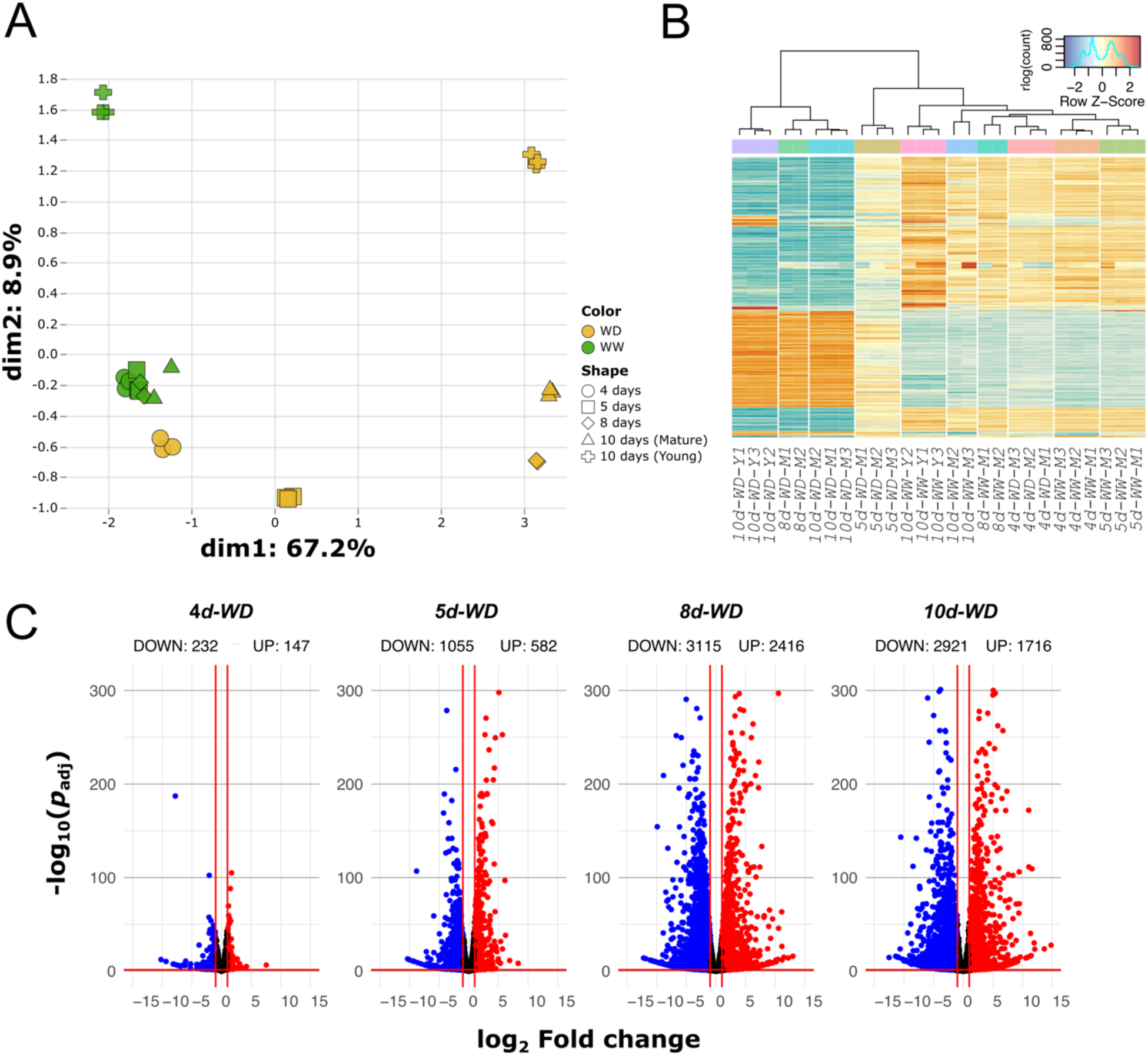
Transcriptomic profiling of water deficit (WD) and well-watered (WW) tomato plants. **A)** Multidimensional scaling plot visualizing (dis)similarity between WD and WW samples. Symbols represent biological replicates, colors represent the different treatments (yellow for WD, green for WW conditions), and different shapes represent the different sampling points (circle for 4d-WD, square for 5d-WD, rhombus for 8d-WD, triangle for 10d- WD, and plus signs for young leaves at 10d-WD). **B)** Expression heatmap and hierarchical clustering of the top 500 most variable genes. Each column corresponds to one replicate: WW or WD, mature (M) or young (Y) leaves, and different clusters are separated by white lines. All biological replicates cluster to their respective time points and treatments. Color is determined by z-score, ranging from -3 (blue) to 3 (red). **C)** Volcano plots showing differentially expressed genes between WW and WD conditions in mature leaves for every time point. Differential expression is defined as having a log_2_ fold change over WW conditions of at least 1, and a Benjamini-Hochberg FDR-adjusted *p*-value below 0.05.

We compared gene expression in the well-watered and drought-stressed plants for each time point. In total, we found 7179 genes that were differentially expressed (DE) in at least one time point (Suppl. Table S2). At 4 days after water deficit (4d-WD at 10% reduction in *gs*), a total of 379 genes were differentially expressed (232 genes were down- and 147 genes were upregulated; Fig. 2C). At 5d-WD at 90% reduction in *gs* (i.e., the stomata were fully closed) we found 1637 DE genes (1055 down- and 582 upregulated; Fig. 2C). At 8d- WD after the leaves had fully wilted and embolism started to develop, we found 5531 DE genes (3115 down- and 2416 upregulated; Fig. 2C). Lastly, at 10d-WD, when the vulnerability curve started to show an exponential increase in embolism events and the percentage of embolized pixels (PEP) reaches 12% (P_12_; Fig. 1B), we found 4637 DE genes (2921 down- and 1716 upregulated; Fig. 2C).

There was a pronounced difference in the number of pre- and post-stomatal closure DE genes (Suppl. Fig. S5). Most DE genes were unique to each time point, highlighting the particular importance of transcription for the specific WD intensity. However, 105 genes were commonly up- or down-regulated during all WD intensities, whereas a much larger portion (1011) was differentially expressed in post-stomatal closure samples (from 5d- to 10d-WD), indicating an important shift in the leaf physiology imposed by (and/or stimulating) stomatal closure. Similarly, a large fraction of DE genes (2514) is shared across the post-wilting samples, when embolism starts to develop in leaf xylem.

Next, we examined the clustering and expression patterns of genes via a weighted gene correlation network analysis (WGCNA, Fig. 3). Firstly, we restricted our analysis only to the previously identified 7179 genes that showed differential expression in at least one time point (Suppl. Table S2). To effectively characterize gene expression under increasing WD, we used a pseudo-time series, taking 4d-WD well-watered plants as basal condition, following up with WD samples for subsequent time points (4-10-d WD). Nineteen clusters of inter-correlated genes were identified (Fig. 3A). We correlated the cluster eigengenes to the physiological traits we measured on the same plants (Fig. 3B), showing their change over increasing WD. Three groups of clusters were easily detectable. The first group (clusters i-vi) showed a positive relationship between gene expression and Ψ_leaf_, *gs*, and *A* (Fig. 3B), a negative relationship with the increase of embolism (leaf PEP, Fig. 3B), and a general downregulation during increasing WD (Fig. 3C). The second group (clusters vii-xiii) showed the opposite trend: a negative relationship between gene expression and Ψ_leaf_, *gs*, and *A* (Fig. 3B), a positive relationship with the increase of embolism (leaf PEP, Fig. 3B), and a general upregulation during WD (Fig. 3C); the third group (clusters xiv-xix) showed an intermediate regulation compared to the two previous groups and a significant relationship with few or no recorded traits.

**Figure 3.**
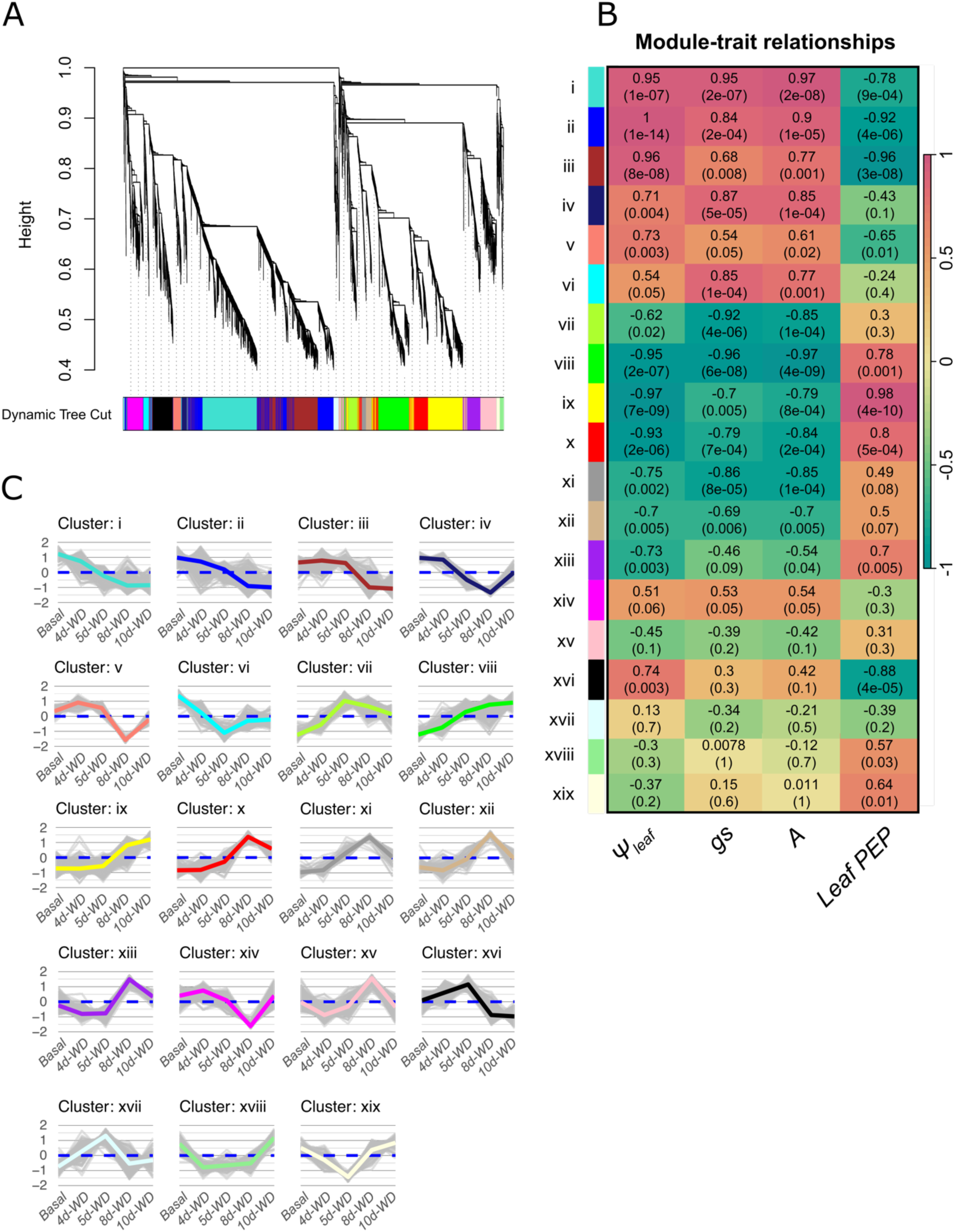
Weighted gene correlation network analysis (WGCNA). **A)** Dendrogram denoting WGCNA results for all previously identified DE genes. The y-axis (Height) represents the distance, or dissimilarity between clusters. Nineteen distinct co-expression clusters were identified, with different colors and Roman numerals (i-xix). Gene cluster designations can be found in Suppl. Table S2. **B)** Cluster-trait relationship heatmap, correlating the module eigengenes (ME) to the measured physiological traits: leaf water potential (Ψ_leaf_), stomatal conductance (*gs*), CO_2_ assimilation (*A*), and loss of hydraulic conductance in leaves (leaf PEP). Correlations are plotted on a diverging color scale centered on 0 (yellow), with red denoting a positive correlation and green a negative correlation. **C)** Expression Z-scores for each cluster, using scaled and centered VST normalized counts over increasing water deficit from basal (WW) conditions, consisting of WW plants on the first sampling date, to 10d-WD.

### Co-expression gene network analyses indicate key regulatory genes involved in the different stages of the water deficit response

To further explore the transcriptional regulation of the WD response, we prioritized identifying the most vital transcription factors (TFs) given their pivotal roles in regulating gene expression. We argued that TFs are important for two primary reasons: 1) they could exhibit the highest differential expression at a given time point and 2) they could exert a high influence on the network. To this end, we calculated centrality measures (degree and betweenness centrality) for all previously identified TFs of interest, in addition to examining differential expression per time point (Fig. 4; Suppl. Fig. S6). Two TFs in particular (*Solyc01g095460.3* and *Solyc05g050220.3*; Suppl. Fig. S6) presented the highest number of correlations and betweenness centrality over the four sampled dates. These two TFs belong to the G-box binding factor (GBF) family and their expression increases with increasing WD (cluster viii). They are close homologs of *AtGBF3*, an abscisic acid (ABA)-activated Arabidopsis gene that confers increased tolerance to drought when overexpressed (Lu *et al.*, 1996; Ramegowda *et al.*, 2017). ABA is a critical hormone in abiotic stress signaling known to accumulate during WD stress (Waadt *et al.*, 2022). Our data suggest that both genes play a pivotal role in the drought response of tomato leaves, likely by triggering a cascade of events related to the ABA-dependent response to WD.

**Figure 4:**
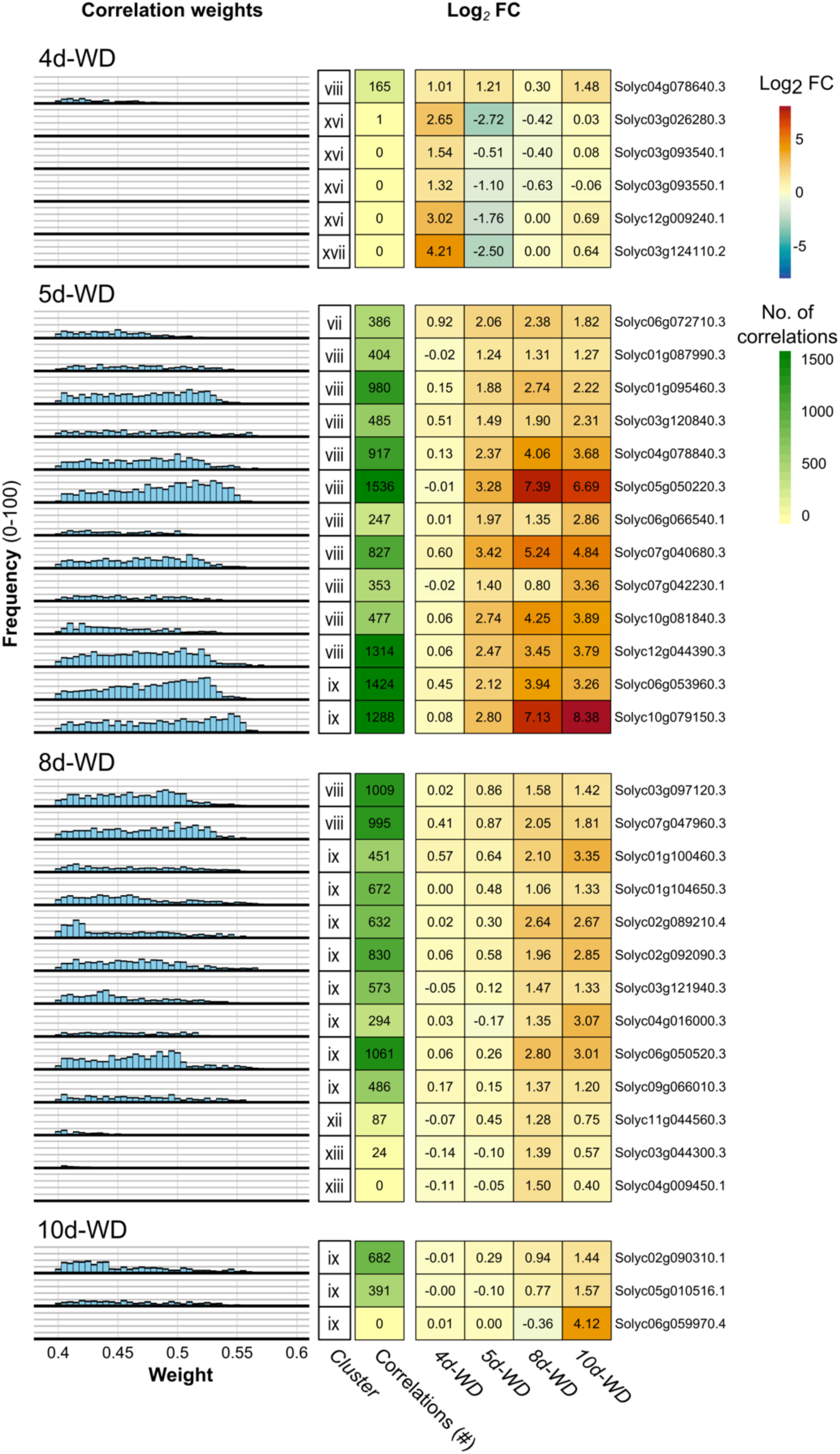
Transcription factor expression and number of correlated genes over a progressive water deficit. Transcription factors are separated by which sampling time point they first break the differential expression threshold (log_2_ fold change > 1; *p_adj_*< 0.05), focusing only on upregulated transcription factors. In the columns (from left to right) we indicated: a histogram showing the distribution of correlation strengths (weight) of correlated genes, grouping weights within 0.01 difference together; the gene cluster (see Fig. 3); the number of correlated genes from 0 (light yellow) to 2500 (green); the differential expression for each transcription factor across time points is given in log_2_ fold change spanning from -8 (blue) to 8 (red) and centered on 0 (white); and the locus ID of the gene.

Subsequently, we looked for TFs that drive the WD response at the specific time points. We subdivided the previously identified TFs of interest by the time point at which they first show significant upregulation (Fig. 4). At 4d-WD, at the onset of stomatal closure (Fig. 1B), the TFs presenting the highest number of correlations (all belonging to cluster xvi) followed an ABA-independent molecular pathway: *Solyc03g124110.2* and *Solyc03g026280.3* are two genes presenting a strong difference in expression compared to control plants (4.21 and 2.65 LFC, respectively); they are close homologs of Arabidopsis *AtDREB1* (one of the key TFs driving the ABA-independent response to drought; Akhtar *et al.*, 2012). Alongside, *Solyc04g078640.3*, *Solyc12g009240.1*, *Solyc03g093540.1*, and *Solyc03g093550.1* belong to the Ethylene Response Factor (ERF) family, members of which are often, but not always, repressed by ABA (Müller and Munné-Bosch, 2015). It has been shown that hydraulic signals (i.e., changes in water potential) regulate stomatal response to WD before ABA accumulates in the stressed leaves imposing a more stable stomatal closure (Tombesi *et al.*, 2015; Huber *et al.*, 2019; Abdalla *et al.*, 2021). Our data corroborates this, as the ABA-independent TFs were driving the co-expression network in tomato leaflets that showed a reduction of 10% in stomatal conductance and a Ψ_leaf_=-0.41 MPa.

At 5d-WD, the time point of full stomata closure, the leaves were not yet wilting, and the onset of embolism did not yet occur in the leaf xylem (Fig. 1). The TFs presenting the highest number of correlations belong to cluster viii (i.e., increasing linearly with WD intensity) and are mostly related to the ABA-dependent pathway (Fig. 4). More specifically, they included *Solyc01g095460.3* and *Solyc05g050220.3*, close homologs of *AtGBF3* described above, and *Solyc04g078840.3* (*AREB1*, ABA-responsive element binding protein 1), one of the most important TFs in the ABA-dependent response to drought in many plant species (Fujita *et al.*, 2005; Barbosa *et al.*, 2012; Pan *et al.*, 2023). We also retrieved *Solyc12g044390.3*, *Solyc06g066540.1*, and *Solyc03g120840.3*, close homologs of *AtDREB3*, known to be involved in abiotic stress tolerance (Islam and Wang, 2009). In addition to the genes involved in the ABA-dependent response to drought, we found two genes involved in the biosynthesis of antioxidant molecules: *Solyc10g079150.3* and *Solyc10g081840.3* are two *SlNF-YA* TFs involved in flavonol biosynthesis (Li *et al.*, 2016; Wang *et al.*, 2021). Furthermore, *Solyc07g040680.3* and *Solyc06g053960.3* encoding the two heat shock proteins *Sl*HsfA2 and *Sl*HsfA6b, respectively (Huang *et al.*, 2016), were highly correlated as well. In summary, this shift in expression occurring at 5d-WD likely indicates an accumulation of ABA in the leaves of (rather sensitive) tomato plants that become more drought-stressed. The flavonol biosynthesis TFs point to the beginning of the antioxidant response, probably limited to the most sensitive leaf mesophyll tissues (Watkins *et al.*, 2017), and the upregulation of heat shock proteins indicates an increase in leaf temperature resulting from fully closed stomata that strongly reduce water loss and leaf cooling.

At 8d-WD, the leaves were wilting, and embolism events started to develop in the leaf xylem (4-6% PEP). We observed an increase in the number of correlations and betweenness centrality of additional heat-shock responsive TFs *Solyc03g097120.3*, *Solyc04g016000.3*, and *Solyc06g050520.3* indicating increasing leaf temperature and the pronounced role of heat shock proteins in severe WD (Fig. 4; Mishkra *et al.*, 2002; Hwang *et al.*, 2012). Three WRKY TFs were also strongly upregulated, probably related to the antioxidant response (see *SlWRKY01*, *SlWRKY24*, and *SlWRKY35* below). In addition, *Solyc01g100460.3*, *Solyc01g104650.3*, and *Solyc02g092090.3* belonging to the bZIP family were highly correlated, probably involved in C and N metabolism reprogramming (Dietrich *et al.*, 2011). Aside from the analysis of TFs, we observed that genes encoding enzymes for the biosynthesis of trehalose (*Solyc02g071590.3*, *Solyc07g006500.3*) and proline (*Solyc06g019170.3*, *Solyc02g068640.3*) were upregulated from 8d-WD onwards. These are considered two of the most important osmoprotectants, meaning that they regulate the internal cell pressure against the dehydrating apoplast (Iturriaga *et al.*, 2009; Szabados and Savoure, 2010). This indicates that, in these later stages of stressful WD conditions, retaining residual water as much as possible is vital for leaf mesophyll cells. Interestingly, enzymes involved in proline degradation (*Solyc02g089620.3*; *Solyc02g089630.3*) were already downregulated at 4d-WD, demonstrating that the osmoprotectant response is one of the earliest molecular responses to WD in tomato plants.

At 10d-WD, embolism in mature leaves reached P_12_, corresponding to the point after which embolism started to spread exponentially in the leaf veins. Fewer TFs were upregulated, among which *Solyc02g090310.1* and Solyc05g010516.1 presented the highest number of correlations (Fig. 4). Only one of them is well described (*Solyc02g090310.1*); its Arabidopsis homolog, *AtDOF1* (Dof zinc finger protein1), is activated by oxidative stress (Yanagisawa, 2002), indicating the strong presence of reactive oxygen species (ROS) molecules in highly stressed leaves.

### The role and timing of four transcription factor families during water deficit in mature tomato leaves

As an alternative approach to disentangle the transcriptome profiles across the different time points during our tomato drought experiment, we identified genes from four key TF families that presented a significantly higher or lower expression than in control plants (Table 1).

**Table 1.**
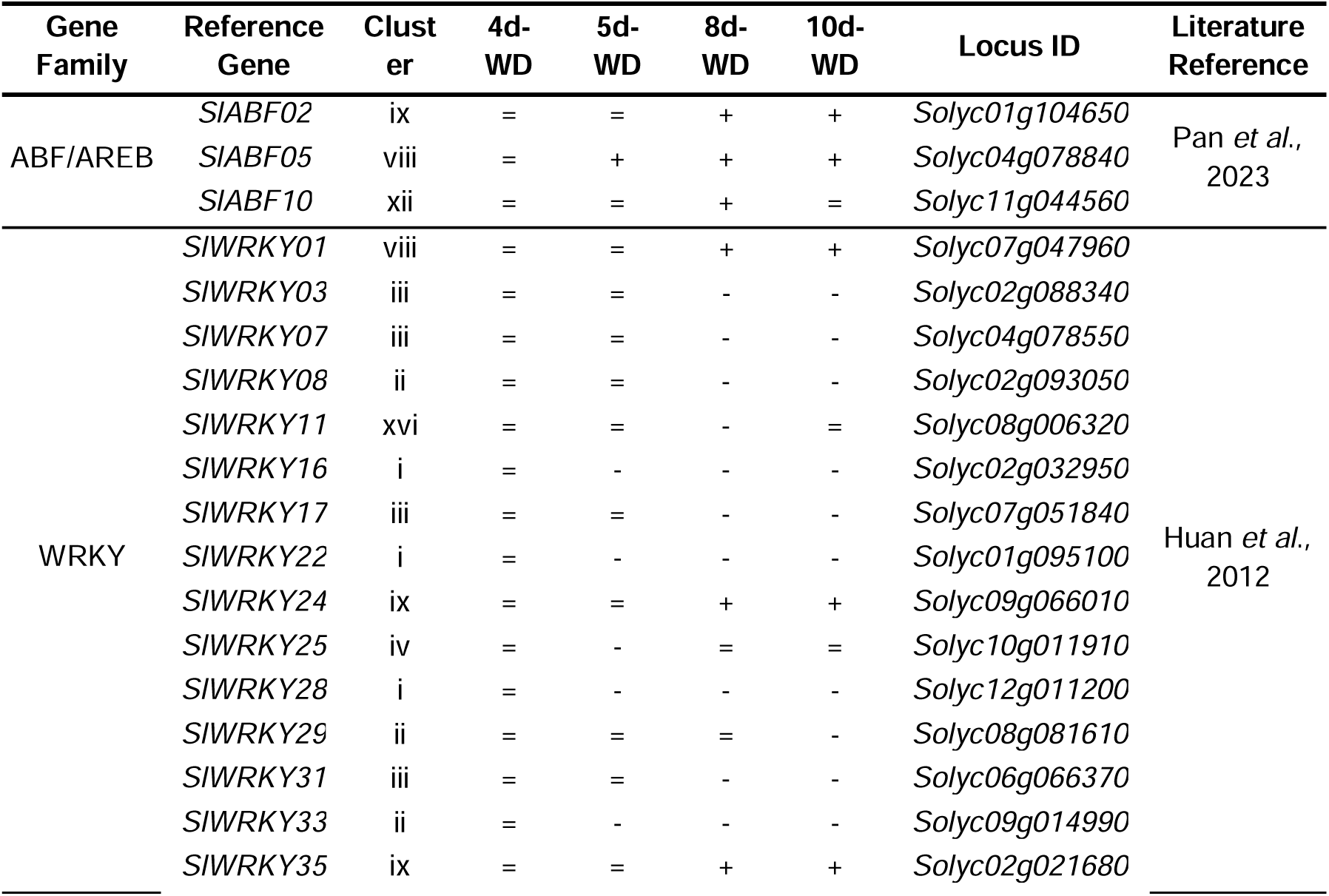
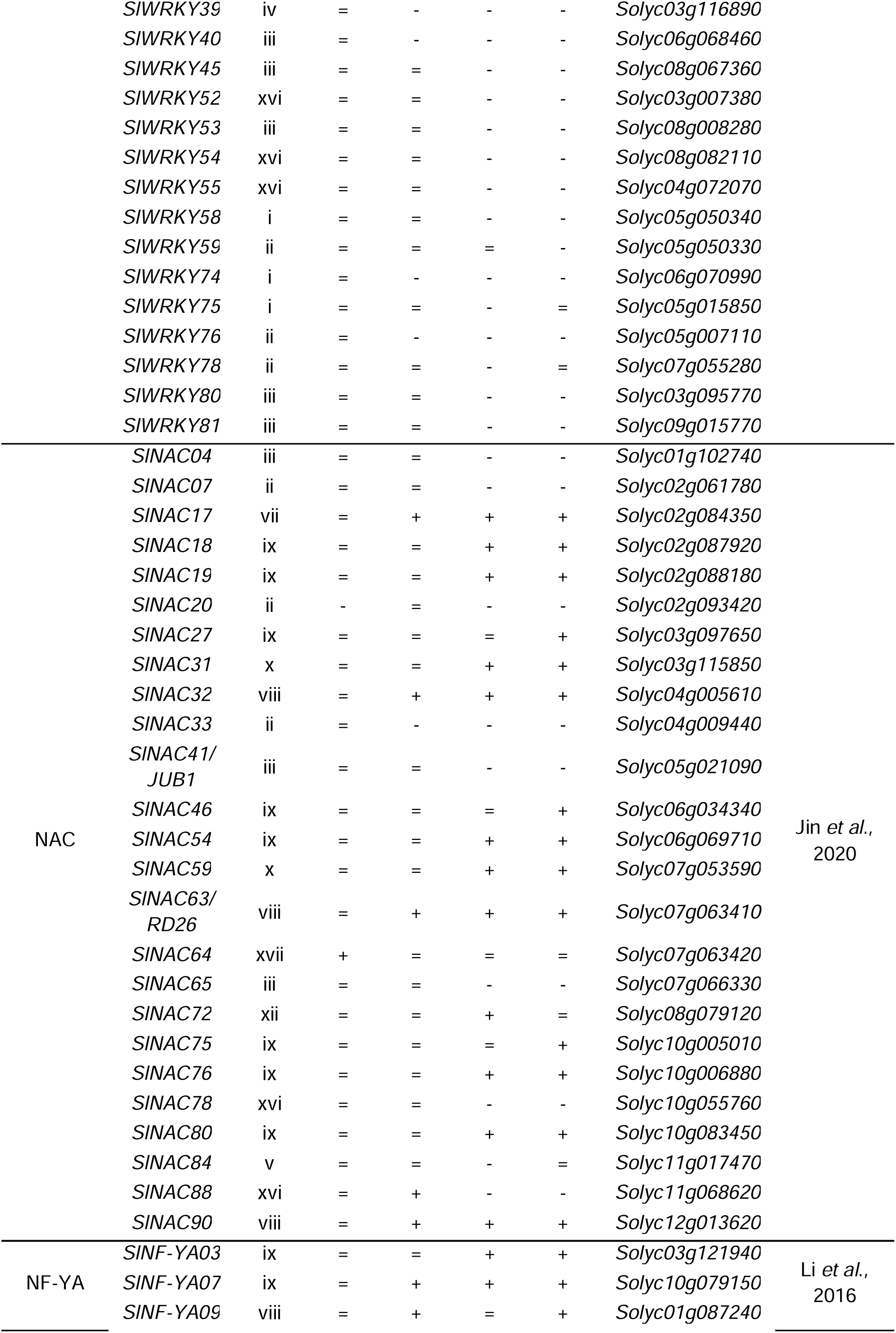
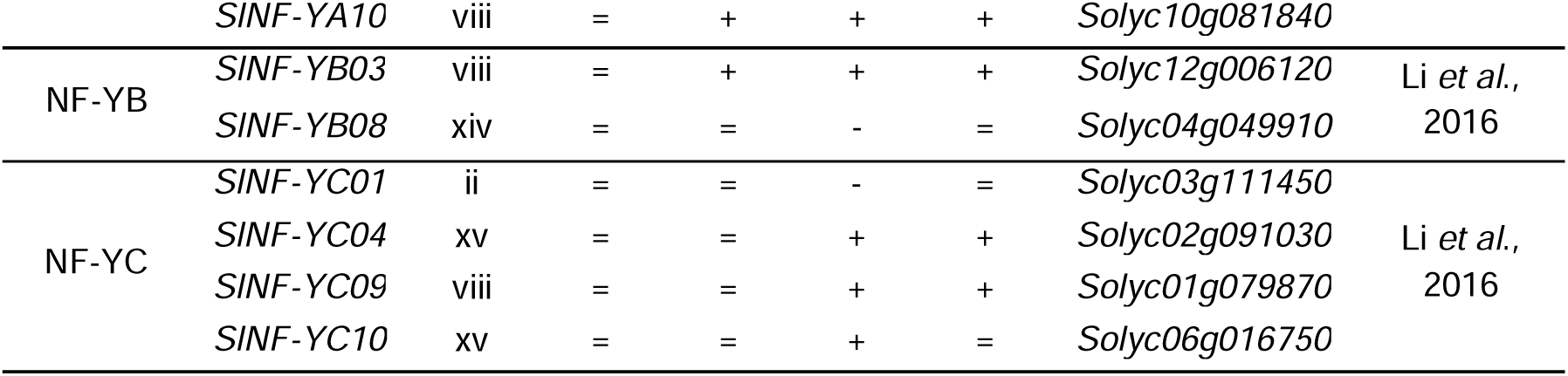
Response of main TF families in tomato along with the genes that were differentially expressed in plants under water deficit in at least one time point compared to control plants (i.e., genes presenting log2 fold change > |1| and *p_adj_*-value < 0.05). For every gene, its cluster of expression (see Fig. 3), its up- (+), down- (-), or no- (=) regulation for different days after drought (4, 5, 8, and 10d-WD) compared to WW plants, and locus ID are indicated, as well as the study describing the function of the different loci.

First, the *ABF*/*AREB* (*ABRE BINDING FACTOR*/*ABA-RESPONSIVE ELEMENT BINDING PROTEIN*) genes encode important transcription factors involved in ABA signaling; they are generally upregulated in the response of plants to abiotic stresses (Pan *et al.*, 2023). We found three genes of this family to be upregulated during WD, confirming their role in the response of tomato to drought (Table 1).

Second, the WRKY family of transcription factors has been described to be involved in plants’ growth development and response to a wide range of abiotic and biotic stresses (Chen *et al.*, 2017). In our experiment, the majority of the differentially regulated *WRKY* genes were found to be downregulated during WD (i.e. 27 genes belonging to clusters i to iv, Table 1), indicating a shift towards *WRKY* deactivation, possibly reducing cell metabolic activities. For example, *SlWRKY81* has been described as a negative regulator of drought tolerance by maintaining open stomata and repressing proline biosynthesis during water shortage (Ahammed *et al.*, 2020a; Ahammed *et al.*, 2020b). In our experiment, expression of *SlWRKY81* was strongly downregulated at the highest intensities of WD. Conversely, three other *WRKY* genes (*SlWRKY01*, *SlWRKY24*, and *SlWRKY35*) previously described to play a role in biotic stress response and carotenoid biosynthesis in tomato (Moghaddam *et al.*, 2019; Yuan *et al.*, 2022), were upregulated during increasing WD. Our results also suggest that these three TFs might be involved in triggering and/or intensifying the antioxidant response of leaves during drought. Four other genes in this family (*SlWRKY11*, *SlWRKY52*, *SlWRKY54*, and *SlWRKY55*) exhibited an intermediate regulation, being upregulated at 5d- WD (at stomatal closure) and downregulated at 8d-WD (leaf wilting and onset of xylem embolism) and 10d-WD (at P_12_; cluster xvi, Table 1). *SlWRKY52* was previously described to have a role during drought stress (Jia *et al.*, 2023), whereby its overexpression generally increased tolerance to drought and osmotic stress in tomato plants. Our results indicate that *SlWRKY52* is activated only during stomatal closure and not at higher WD intensities, confirming the hypothesis of Jia *et al.* (2023) of its potential role in ABA biosynthesis and signal transmission.

Third, the NAC transcription factor family included 7 genes that were downregulated with increasing WD intensity, while 15 genes were upregulated during WD (Table 1). Of the latter ones, *SlNAC63*/*RD26* and *SlNAC90* were previously described to be upregulated during WD (Iovieno *et al.*, 2016), confirming our results showing an expression that increased linearly with increasing stress. Interestingly, three other *NAC* genes (*SlNAC64*, *SlNAC78*, *SlNAC88*) were upregulated only at 5d-WD (Table 1), but downregulated at the other WD intensities, suggesting a specific role during the earlier WD stages at stomatal closure.

Fourth, the nuclear transcription factor Y (NF-Y) family encodes genes that operate in a multiprotein complex consisting of three subunits (NF-YA, NF-YB, and NF-YC). While these TFs are mostly described for their role in flavonol biosynthesis during fruit ripening in tomato (Li *et al.*, 2016; Wang *et al.*, 2021), they were observed to be involved in growth and flower development in multiple plant species (Petroni *et al.*, 2012). Interestingly, *SlNF-YA9*, *SlNF- YB8*, *SlNF-YC1*, and *SlNF-YC9* showed a progressive increase along the increasing WD stages (Table 1), indicating a possible role of these TFs in the antioxidant response in stressed leaves. Likewise, we also found *SlNF-YA10*, which was shown to be involved in increased tolerance to oxidative stress in leaves and fruits of tomato plants (Chen *et al.*, 2020), to be upregulated during increasing WD. Finally, two more NF-Y TFs (*SlNF-YC04* and *SlNF-YC10*) were upregulated exclusively at 8d-WD, the time point of the onset of xylem embolism in the leaves.

### Comparative transcriptomic analyses explain stronger water deficit tolerance of young compared to mature leaves

Young developing leaves had a more negative water potential than mature and fully expanded leaves, both in WW (Ψ_leaf_=-0.5 ± 0.05 MPa versus -0.44 ± 0.07 MPa), and in WD (Ψ_leaf_=-1.6 ± 0.04 MPa versus -1.4 ± 0.07 MPa) conditions. This observation is consistent with the cohesion-tension theory, where an increasingly negative water potential pulls water from the bottom to the top of the plants (Dixon and Joly, 1895). Interestingly, despite this more negative Ψ_leaf_, the young developing leaves at 10d-WD did not show any sign of wilting and/or yellowing, while the mature leaves were well below their wilting point (Suppl. Fig. S1). From an evolutionary standpoint, natural selection may lead to prioritizing the survival of younger leaves over older ones as older leaves are often less efficient in photosynthesis due to age-related decline (Procházková & Wilhelmová, 2007; Wang *et al.*, 2014). This idea is also corroborated by studies highlighting the allocation of resources to younger tissues during stress- and season-related leaf senescence (Schippers *et al.*, 2015; Berens *et al.*, 2019). Likewise, it has been shown that leaves produced later in the season have an increased WD tolerance compared to leaves produced in the early spring in several woody species (Sorek *et al.*, 2021; Sorek *et al.*, 2022).

We aimed to elucidate the mechanisms behind this apparent resilience of younger leaves by comparing the transcriptomic profiles between young and mature leaves under WW and 10d-WD conditions in the same plants. We identified 360 differentially regulated genes (*i.e.*, genes with an LFC (log_2_ fold change) above 2, or below -2, and an FDR-adjusted *p*-value below 0.05) from the interaction between differentially expressed genes between leaf age and watering regime (Suppl. Table S3). We grouped these genes into clusters of similar patterns of expression via K-means clustering. We identified seven distinct gene expression clusters (Fig. 5), which potentially delineate a functional separation in the response to WD. To avoid confusion with our previous clustering related to the WD response in mature leaves only, we named these clusters 1 to 7. To further dive into the potential biological significance of these expression patterns, we performed a Gene Ontology (GO)-enrichment analysis, a manual examination of the *AHRD* gene descriptions, as well as an examination of genes commonly found to be involved in the WD response (Kido *et al.*, 2019; Zhang *et al.*, 2022; Mishra *et al.*, 2023). Below, we limit our discussion to the potential role of the genes in cluster 1 (upregulation of 26 genes in young leaves at 10d-WD) and cluster 5 (downregulation of 97 genes in young leaves at 10d-WD; Fig. 5).

**Figure 5:**
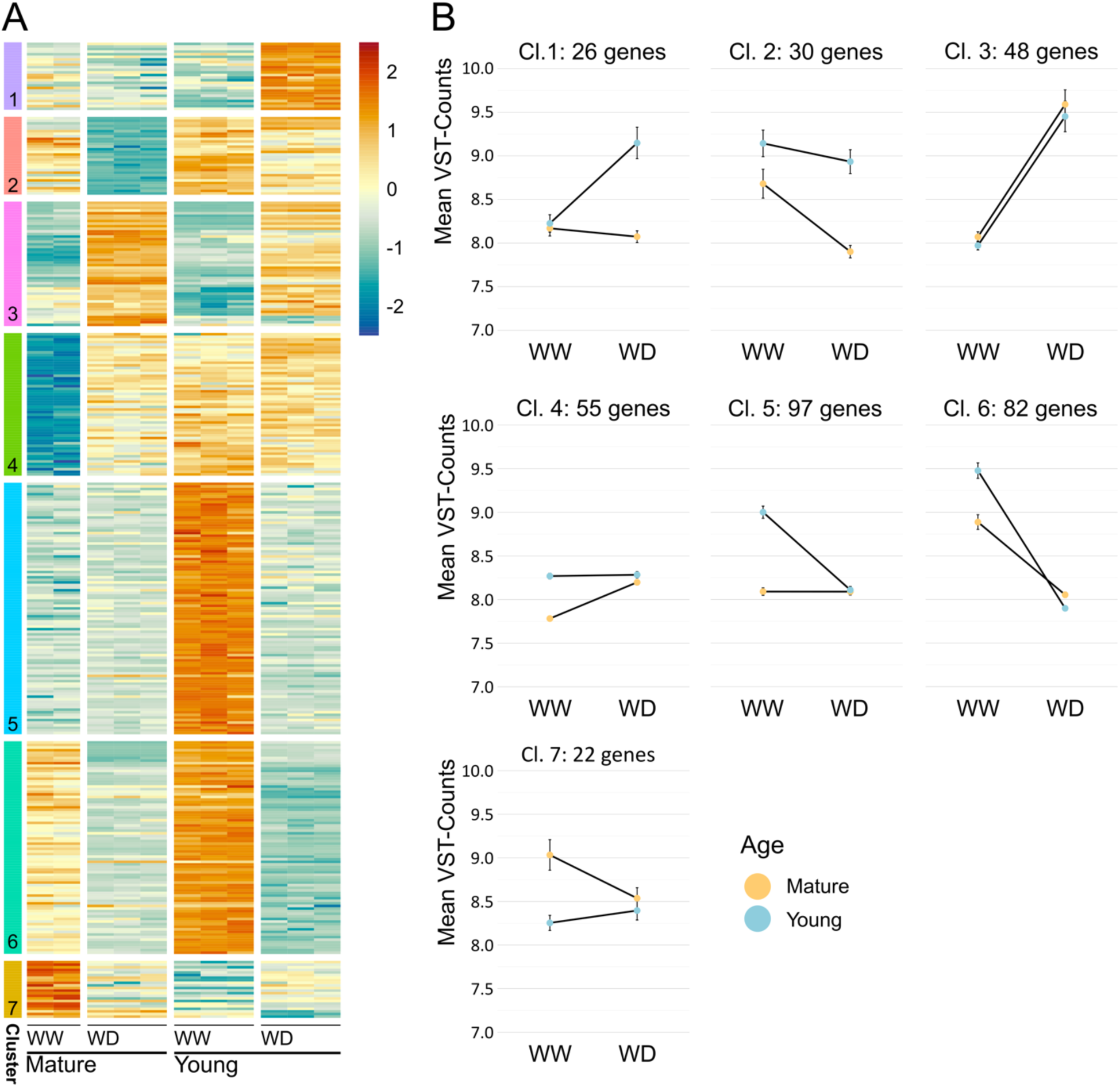
Comparative transcriptomic analysis of young and mature leaves under well-watered and water-deficit conditions. **A)** Heatmap showing z-scores of differentially regulated genes between young and mature leaves under well-watered (WW) and water deficit (WD) conditions (10d-WD). Values are represented by a diverging color scale from low (blue) to high (red), centered on 0 (light yellow). Clusters were determined by K-means clustering and cluster assignments are given on the left. Gene cluster assignments are given in Suppl. Table S3. **B)** Line plots detailing average (± SE) gene expression in each cluster for young and mature leaves under WW and WD conditions. Values are given in Variance Stabilized Transformed (VST) counts. The average gene expression for mature leaves is shown in yellow, and for young leaves in blue.

Cluster 1 contains genes that show an upregulation in WD-young leaves, while they are neither responsive in mature leaves (WW and WD) nor in WW-young leaves (Fig. 5). Thus, this cluster could potentially aid in conferring a WD tolerance specifically to younger leaves. For instance, we found that cluster 1 is enriched for genes involved in cell wall biosynthesis (Suppl. Fig. S7) with multiple xyloglucan endotransglucosylase enzymes (XTH; *Solyc03g093120.5, Solyc03g093080.3, Solyc03g093130.3*). XTHs catalyze the cleavage and (often) re-ligation of xyloglucan molecules to cellulose and can have dual roles in cell wall rigidity (Kaku *et al.*, 2004; Miedes *et al.*, 2013). In Arabidopsis, constitutive overexpression of hot pepper (*Capsicum annuum*) *CaXTH3* enhanced tolerance to salinity and WD, likely through increased cross-linking of xyloglucan molecules, thereby decreasing plasticity and elongation of the cell walls (Cho *et al.*, 2006; Hayashi & Kaida, 2011). It is plausible that increased cell wall deposition and rigidity in younger leaves contribute to the maintenance of turgor pressure in smaller cells, at the expense of cell division and expansion (i.e., ceased leaf growth). This hypothesis is corroborated by the increased expression in young leaves of cellulose synthase (*Solyc02g072240.3*) and fasciclin-like arabinogalactan protein (*Solyc11g069250.2*) in cluster 4, and the decreased expression of expansins (*Solyc09g018020.3, Solyc06g049050.3*) in cluster 5.

Cluster 1 also contains a callose synthase (*Solyc11g005980.3*). Callose deposition is intricately connected to plasmodesmata permeability and is tightly controlled by two antagonistic enzymes: the synthesizing callose synthases/glucan synthase-like protein (CS/GSL), and breakdown β-1,3-glucanases (BGs) (Wu *et al.*, 2018). Interestingly, one of the closest homologs of *Solyc11g005980.3* in Arabidopsis (*AtGSL12*/*AtCalS3*) has been connected to callose deposition in sieve elements in the phloem, where increased activity led to clogging of the plasmodesmata pores and impaired phloem loading (Vatén *et al.*, 2011; Wu *et al.*, 2018). In our dataset, one BG (*Solyc01g060020.4*) is strongly downregulated during WD uniquely in young leaves. The increased expression of the callose synthase gene *Solyc11g005980.3*, together with the decreased expression of the callose breakdown gene *Solyc01g060020.4* in young tomato leaves during WD conditions suggests a combined approach to stimulate callose deposition in the phloem thereby blocking outward sugar transport. Soluble sugar accumulation during WD has been related to an increased osmoprotective power delaying leaf wilting (Ozturk *et al.*, 2020), as well as a ready source of energy during the antioxidant response (Sami *et al.*, 2016). Rosa *et al.* (2009) highlighted a whole chain of gene up- and down-regulations coordinated by soluble sugars during abiotic stresses. Our transcriptomic profile also suggests that young leaves accumulate sugars, which in turn might be related to (if not coordinating) the accelerated response to dehydration in these developing leaves.

Cluster 5 contains genes that show a much higher expression in young leaves under WW conditions, while they are downregulated to the same level as in mature leaves during WD (Fig. 5). This cluster may contain genes that are “shut off” during WD in younger leaves. Aside from the previously mentioned expansins, this cluster is enriched with nine genes encoding for chlorophyll a/b binding proteins (*Solyc08g067330.1, Solyc02g070980.1, Solyc06g069730.3, Solyc02g070970.1, Solyc02g070990.1, Solyc02g071000.1, Solyc02g070950.1,* and *Solyc02g071010.1*). The increased expression of the chlorophyll a/b binding protein genes under WW conditions is indicative of leaves undergoing active growth, as actively proliferating cells build their photosynthetic machinery (Cackett *et al.*, 2022). The reduction in expression of these genes during WD coincided with the shifting in cell wall structure (and consequently growth cessation) highlighted in cluster 1, indicating that photosynthesis as well is affected at this WD intensity. Moreover, even under well-watered conditions, the photosynthetic apparatus is one of the major producers of ROS, which is exacerbated by abiotic stress (Asada *et al.*, 2006; Gill & Tuteja, 2010). Therefore, the capacity of young leaves to more effectively reduce photosynthetic activity compared to older leaves may also help to reduce the oxidative pressure under WD.

We also found that young leaves maintain an overall higher presence of negative regulators of peptidase activity under WD. Two are upregulated uniquely in young leaves under WD (cluster 1; *Solyc09g083435.1, Solyc03g098795.1*) and three remain highly expressed in young leaves, while they are downregulated in mature leaves (cluster 2), namely one cathepsin D inhibitor protein (*Solyc03g098790.3*), and two protease inhibitors (*Solyc09g083435.1, Solyc03g098795.1*). In addition, eleven prote(in)ase inhibitors maintained a significantly higher expression only in WD-young leaves, compared to just three which maintained a higher expression in WD-mature leaves. Proteases are incredibly varied, and their functions may be both detrimental and beneficial for acquiring tolerance to WD (Jedmowski *et al.*, 2014; Cheng *et al.*, 2016; Malefo *et al.*, 2020; D’Ippólito *et al.*, 2021; Moloi & Ngara, 2023). The increased presence of prote(in)ase inhibitors activity in young leaves under WD could denote decreased protease activity. Although increased protease activity may aid in maintaining a functional protein/enzyme pool in the face of oxidative stress by increasing protein turnover, having a high protein turnover is resource-intensive and may not be sustainable in a prolonged drought. Considering the time point of measurement (10d-WD) it would be plausible that there is a focus on maintaining current proteins, specifically in young leaves.

## Conclusions

By disentangling gene expression profiles at precise physiological thresholds during increasing water deficit in mature leaves, we identified key regulatory genes and main TF families that play a role in the drought response of tomato leaves. We highlighted the activation of ABA-independent TFs during the lowest WD intensities (at 4d-WD with partial stomatal closure, i.e. 10% reduction in stomatal conductance (*gs*)), followed by a high transcription of ABA-dependent TFs one day later (5d-WD, full stomatal closure, i.e. 90% reduction in *gs*). At stronger WD intensities after 8 days, when embolism started to develop in the leaves (4-6% PEP), we observed an activation of *HEAT SHOCK PROTEIN* genes and a general antioxidant response. Finally, at the strongest intensities after 10 days of WD, when embolism spread is about to exponentially increase (P_12_), TFs related to oxidative stress were highly upregulated. When transcriptome profiles of young versus mature leaves are compared after 10d-WD in the same tomato plants, we can generalize that the stronger resilience of younger leaves to WD results from a shift in resource allocation from cell proliferation and expansion toward homeostasis maintenance. Compared to mature leaves, young leaves showed increased expression of genes altering the cell wall structure, like XTHs, cellulose synthase, and fasciclin-like arabinogalactan genes, which possibly create smaller cells with more rigid cell walls that can sustain stronger turgor pressures. Moreover, a potential preferential callose deposition at the phloem sieve elements could minimize water, sugar, and nutrient loss from these young leaves, allowing them to maintain osmoprotectant capacity and energy storage. Finally, by downregulating chlorophyll a/b binding protein genes, young leaves might reduce photosynthesis-related ROS production. This combined ecophysiological-molecular dataset in tomato leaves illustrates how gene regulatory pathways and ecophysiological thresholds are tightly intertwined. We anticipate that these outcomes will serve future studies aiming to integrate both disciplines more thoroughly, which will undoubtedly shed new light on how natural and agricultural selection has impacted the underlying mechanisms of drought response across the plant kingdom.

## Supplementary information

**Supplementary Figure S1**. Visual change of plant morphology over increasing water deficit.

**Supplementary Figure S2**. A compound leaf from the 6-8th node, showing leaflets selected for gas exchange, stomatal size, RNA-Seq, leaf PEP, Ψ_TLP,_ and water potential analyses.

**Supplementary Figure S3**: Clustering accuracy as viewed on the first three principal components.

**Supplementary Figure S4**. Scale-free topology model fit and mean connectivity for different soft threshold powers.

**Supplemental Figure S5:** Venn diagram of up- and downregulated genes per time point.

**Supplementary Figure S6:** Transcription factors co-expression network.

**Supplementary Figure S7:** Gene ontology enrichment.

**Supplementary Table S1:** Variance Stabilizing Transformation (VST) Counts.

**Supplementary Table S2:** Differentially Expressed (DE) genes in mature leaves.

**Supplementary Table S3:** Differentially Expressed (DE) genes in young leaves at 10 d-WD.

## Acknowledgments

We would like to thank Alex Bos (IBL), Jan Vink (IBL), Bertie-Joan van Hueven (Naturalis), Rob Langelaan (Naturalis), and Régis Burlett (platform Phenobois Bordeaux) for providing technical help. We want to thank Dirk Walther (MPI-MP) for verifying the bioinformatic approaches. B.M.-R. and T.d.W. thank the International Max Planck Research School ‘Primary Metabolism and Plant Growth’ (IMPRS-PMPG; fellowship to T.d.W.) for support. F.L., M.L., and G.B. thank the Dutch Research Council for funding (grant ALWOP.488).

## Author contributions

G.B., T.d.W., B.M.R., S.B., and F.L. designed the experiments. S.B., F.L., and B.M.-R. initiated the research and provided funding. G.B., M.L., and A.T. carried out the physiological measurements. T.d.W. carried out the differential expression and clustering analyses. G.B. and T.d.W. examined genes of interest, identified structural homologs in Arabidopsis, and performed accompanying literature research. G.B. and T.d.W. wrote the manuscript by accepting amendments from all authors. All authors agreed on the final version of the manuscript.

## Data availability

Raw aligned and mapped counts from all samples, as well as the raw sequencing files are available at https://www.ncbi.nlm.nih.gov/geo/query/acc.cgi?acc=GSE270245. Differential expression results, as well as the VST-stabilized counts as used for the WGCNA network generation are provided in the Supplementary Data.

## Figures alt text

Figure 1A: Line graph showing leaf water potential variation over time, for both well-watered (green) and water-deficit subjected (yellow) tomato plants during a 10-day drought experiment.

Figure 1B: Graph showing stomatal safety margin during leaf water potential decline. The blue curve (right) shows gas exchange, the red curve (left) shows the percentage of embolized pixels. A purple vertical line marks the turgor loss point.

Figure 2A: Multidimensional scaling plot showing dissimilarity between water deficit (yellow symbols) and well-watered (green symbols) tomato plant samples. Shapes represent sampling points.

Figure 2B: Heatmap and hierarchical clustering of the top 500 most variable genes in water deficit and well-watered tomato plants.

Figure 2C: Volcano plots showing differentially expressed genes between well-watered and water deficit conditions in mature leaves at each time point. Differential expression is defined as log2 fold change >1 and FDR-adjusted p-value <0.05.

Figure 3A: Dendrogram showing WGCNA results for differentially expressed genes in tomato plants under water deficit. 19 distinct co-expression clusters are identified with different colors and Roman numerals.

Figure 3B: Heatmap showing cluster-trait relationships between module eigengenes and physiological traits: leaf water potential (Ψleaf), stomatal conductance (gs), CO2 assimilation (A), and leaf PEP.

Figure 3C: Line plot showing expression Z-scores for each WGCNA cluster over increasing water deficit from 0 to 10 days. Expression values are scaled and centered on VST normalized counts.

Figure 4: Heatmap showing transcription factor expression and number of correlated genes over progressive water deficit in tomato plants. Transcription factors are separated by the time point they first show differential expression.

FIgure 5A: Heatmap showing z-scores of differentially regulated genes between young and mature leaves under well-watered and water deficit conditions at 10 days.

Figure 5B: Line plots showing average gene expression (± SE) in each K-means cluster for young and mature leaves under well-watered and water deficit conditions.

**Supplementary Figure S1:** Series of images (A-E) showing visual changes in tomato plant morphology over increasing water deficit from well-watered to 10 days after water deficit. Image F shows a sampled mature compound leaf and three younger leaves from the same plant at 10 days.

**Supplementary Figure S2:** Image of a compound leaf from the 6-8th node, highlighting leaflets selected for various analyses.

**Supplementary Figure S3:** Scatter plot showing clustering accuracy on the first three principal components.

**Supplemental Figure S4:** Two-panel graph showing scale-free topology model fit and mean connectivity for different soft threshold powers. The left panel shows model fit (y-axis) against soft threshold powers (x-axis), and the right panel shows mean connectivity (y-axis) across the same range of powers.

**Supplemental Figure S5:** Venn diagram showing numbers of up- and downregulated genes at each time point and their intersections. Numbers in sections represent gene counts, with percentages indicating the fraction of total differentially expressed genes.

**Supplemental Figure S6:** Network diagram showing transcription factor co-expression network in tomato plants under well-watered and water deficit conditions.

**Supplemental Figure S7:** Dot plot showing significantly enriched gene ontology terms on biological processes within four clusters.

